# mRNA delivery of dimeric human IgA protects mucosal tissues from bacterial infection

**DOI:** 10.1101/2023.01.03.521487

**Authors:** Cailin E. Deal, Angelene F. Richards, Tracy Yeung, Max J. Maron, Ziqiu Wang, Yen-Ting Lai, Brian R. Fritz, Sunny Himansu, Elisabeth Narayanan, Ding Liu, Rositsa Koleva, Stuart Licht, Chiaowen J. Hsiao, Ivana L. Rajlic, Hillary Koch, Michael Kleyman, Mark E. Pulse, William J. Weiss, Jennifer E. Doering, Samantha K. Lindberg, Nicholas J. Mantis, Andrea Carfi, Obadiah J. Plante

**Affiliations:** Moderna, Inc.; Cambridge, Massachusetts, USA; HSC College of Pharmacy, University of North Texas; Fort Worth, Texas, USA; Division of Infectious Diseases, Wadsworth Center, New York State Department of Health; Albany, New York, USA; Department of Biomedical Sciences, University at Albany School of Public Health; Albany, New York, USA

**Keywords:** mRNA, antibody, mucosal, infection, therapy

## Abstract

Monoclonal antibody (mAb) therapy is a promising infectious disease intervention strategy but is limited to IgG1 isotypes that have restricted access to mucosal sites. IgA is well-established as the predominant antibody isotype in mucosal secretions but is clinically underutilized. To enable development of IgA-based mAbs, we exploited mRNA platform technology and demonstrated expression of functional, antigen-specific IgA (IgA_mRNA_) that can limit bacterial invasion in the intestine and prevent colonization in the lung. Moreover, *in vivo* IgA_mRNA_ had enhanced serum half-life and a greater degree of sialylation than a recombinantly produced IgA. The results underscore the potential of mRNA-based platforms to deliver protective human mAbs to mucosal surfaces and open new avenues to combat infectious diseases in the face of pervasive antibiotic resistance.

**One Sentence Summary:** mRNA-encoded human monoclonal IgA traffics to mucosal tissues and provides protection against bacterial challenge

## Main Text

Immunoglobulin A (IgA) is the predominant antibody isotype in mucosal secretions of the gastrointestinal (GI) tract and upper airways, where it functions as a first line of defense against invading pathogens (*1, 2*). Humans have two IgA subclasses, IgA1 and IgA2, that differ in hinge length, degrees of O-linked glycosylation (*1*), and mucosal localization. While serum IgA is typically monomeric (_m_IgA), mucosal IgA is dimeric (_d_IgA) or polymeric due to the presence of the joining chain (JC) that covalently links two or more IgA monomers together via their C-terminus (*3*). Human _d_IgA (but not _m_IgA) is transported across epithelial cell barriers of the gut and airways by the polymeric immunoglobulin receptor (pIgR) resulting in secretion of _d_IgA that functions in entrapment of pathogens and prevents attachment to epithelial receptors at mucosal surfaces.

For reasons related to ease of manufacturing and favorable pharmacokinetic properties, immunoglobulin g (IgG) has been the isotype of choice for monoclonal antibodies (mAbs) (*4*). Recombinant IgA (IgA_R_), in contrast, has proven challenging on several fronts. Human IgA, for example, is heavily glycosylated, which raises concerns from a biopharmaceutical standpoint as N-glycosylation can influence conformation, thermal stability, folding efficiency, solubility, and susceptibility to proteolytic degradation (*5-8*). Moreover, IgA_R_ is intractable for most clinical purposes, as it exhibits a shorter serum half-life than endogenous human IgA due to the clearance from circulation as a result of incomplete sialylation (*9*) and inability to recycle through the neonatal Fc receptor (*10*). Efforts to generate IgG/IgA chimeras with desired Fc receptor interactions, serum half-life, and mucosal delivery are ongoing (*11-13*) but have not reached clinical stage readiness.

The severe acute respiratory syndrome coronavirus 2 (SARS-CoV-2) pandemic has highlighted the speed and untapped potential of mRNA technology (*14*). Fundamental to the mRNA platform is the ability to generate difficult-to-manufacture protein complexes *in vivo*, and bypass traditional protein manufacturing and purification (*15*). Though most of the attention for mRNA technology has been on vaccines, the potential use for expression of human proteins with therapeutic potential is expanding. Recent peer-reviewed studies have reported on the delivery and characterization of mRNA-based IgG mAbs in mice (*16*) as well as the safety and efficacy of mRNA delivery IgG mAbs in a phase I clinical trial (*17*), although none to our knowledge have examined the potential to deliver human IgA as a tool to prevent mucosal infections.

In this report, we investigated the use of mRNA for *in vivo* production of structurally and functionally intact human IgA. As proof of principle, we employed two well-characterized mAbs that have proven activity in extensively characterized mouse models of mucosal challenge. Sal4 is an IgA mAb that recognizes the O5-antigen of lipopolysaccharide (LPS) from *Salmonella enterica* serovar Typhimurium (STm) (*18*), and has been shown to reduce invasion of STm into intestinal Peyer’s patches when administered passively (*18, 19*). CAM003 is an IgG1 mAb that binds to *Pseudomonas aeruginosa* (PA) biofilm component Psl and has demonstrated protection in multiple different PA animal models, including acute lung pneumonia (*20, 21*). Here, Sal4 and CAM003 were encoded as mRNA as a human IgA2 (IgA2_mRNA_) or a human IgA1 (IgA1_mRNA_), respectively, and were characterized *in vitro* and *in vivo* as compared to analogous recombinant proteins (IgA2_R_) to assess the potential of the mRNA platform to generate potent and protective mucosal mAbs.

## Results

### Production of functional IgA2 from mRNA is indistinguishable from IgA2_R_ in biophysical and functional assays in vitro but displays better pharmacokinetic properties in vivo

To explore the potential of mRNA to encode functional monoclonal _d_IgA (*18, 19*), mRNA encoding Sal4 heavy chain (HC) and light chain (LC) were packaged in lipid nanoparticles (LNPs), with or without mRNA encoding JC. Transfection of HEK293 cells resulted in IgA2_mRNA_ antibody that recognized STm by enzyme-linked immunosorbent assay (ELISA) to a level similar to IgA2_R_ (**Fig 1A**). Only Sal4 IgA2_mRNA_ transfected with JC were bound by immobilized pIgR, indicating the formation of _d_IgA (**Fig. 1B**). This was validated by chromatography, where there was a predominance of _d_IgA2 over _m_IgA2 upon JC coexpression, as compared to _m_IgA2 or aggregates of IgA2 that assembled in the absence of the JC (**Fig. 1C**) (*22*). Notably, size exclusion chromatography retention times of the _d_IgA2 and _m_IgA2 peaks were nearly identical between IgA2_mRNA_ and IgA2_R_ (**Fig. S1A**). Negative stain electron microscopy (EM) images of the _d_IgA2_mRNA_ preparations revealed images of immunoglobulins in a tail-to-tail oligomerization configuration identical to that of _d_IgA2_R_, demonstrating the ability of mRNA to encode for oligomeric mAbs (**Fig. 1D; Fig. S1B; Fig. S2**). Functionality of IgA2_mRNA_ was assessed in an *in vitro* assay of STm HeLa cell invasion. mRNA transfections (with JC) of EXPI293 resulted in functional IgA2_mRNA_ that reduced STm invasion of HeLa cells to a level similar to IgA2_R_ (**Fig. 1E**) (*19*), demonstrating for the first time that mRNA can encode for functional oligomeric IgA proteins.

**Fig. 1.**
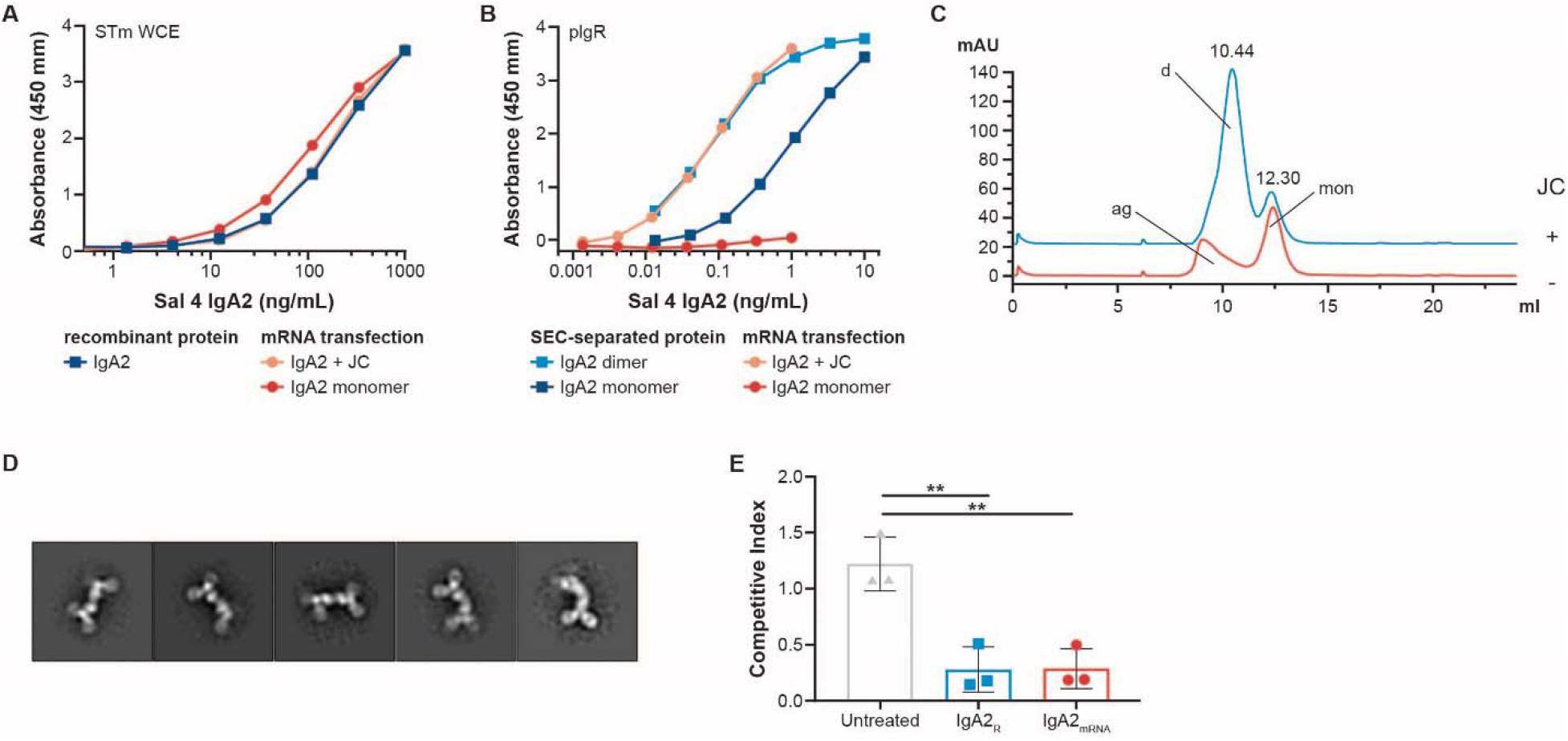
Characterization of Sal4 IgA2_mRNA_ from in vitro transfection. **(A)** Binding of mRNA transfection supernatant and recombinant protein of Sal4 IgA2. **(B)** Binding of SEC-separated Sal4 IgA2_R_ and supernatant from mRNA transfected Sal4 IgA2 to immobilized human pIgR. **(C)** Analytical size exclusion chromatogram of affinity purified IgA2 from mRNA transient transfection. Dimer (d) and monomer (mon) peaks are denoted. Transfections with JC are in blue and without JC are in orange with aggregates labeled (ag). **(D)** Representative image of negative stain EM 2D class averages of the dimeric peak isolated from mRNA transfection with J chain. **(E)** Effect of IgA2_R_ or IgA2_mRNA_ on STm invasion of HeLa cells. ** P<0.05 as determined by unpaired Student’s t test (n=3 experiments done in triplicate).

While the mucosal targeting and oligomeric properties of IgA are clinically appealing, these proteins have traditionally suffered from poor pharmacodynamic properties and in particular, a short serum half-life (*9, 23*). To assess how *in vivo* production of IgA from formulated mRNA (containing HC, LC, and JC) impacts pharmacodynamics, BALB/c mice were injected with 5mg/kg IgA2_R_ (containing a mixture of monomeric and dimeric), 4.5 mg/kg polyclonal human IgA isolate from serum (mostly monomeric) and mRNA/LNP encoding IgA2_mRNA_. Similar to previous reports (*23*), IgA2_R_ exhibited a terminal half-life of 0.64 days and was detected for 2 days while polyclonal human serum IgA was detected for 12 days with a terminal half-life of 0.93 days (**Fig. 2A**). Excitingly, IgA2_mRNA_ exhibited a terminal half-life of 1.63 days, demonstrating a fundamental divergence from that of IgA2_R_ (**Fig. 2A, B**).

**Fig. 2.**
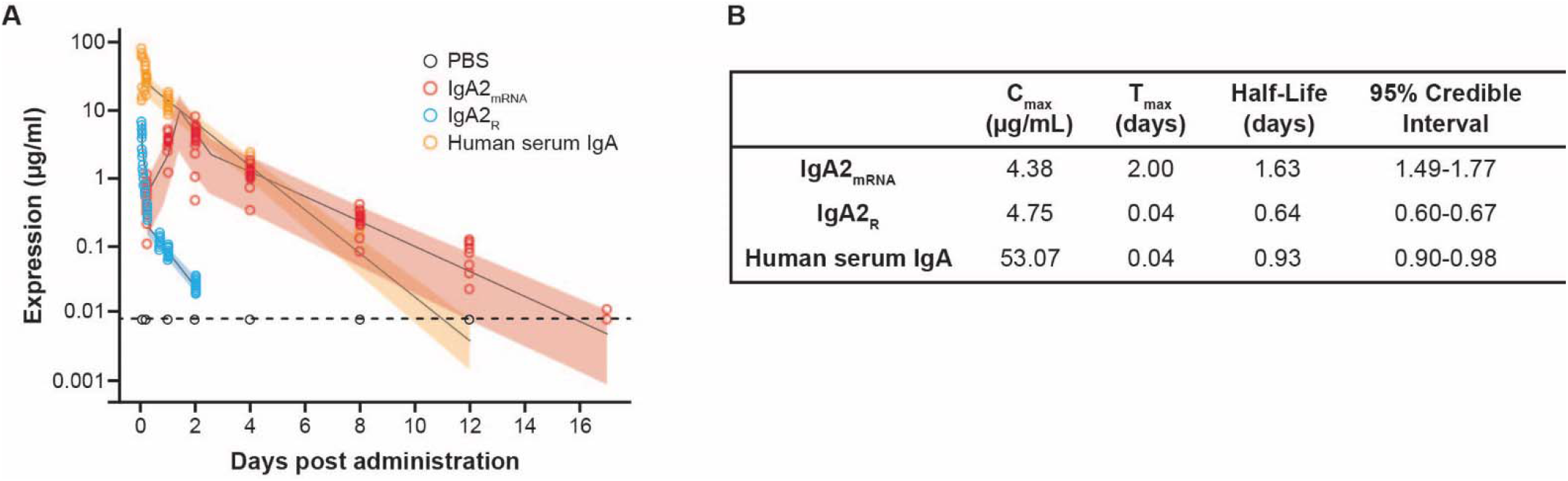
Characterization of IgA2_mRNA_ *in vivo*. BALB/c mice were injected intravenously with 1 mg/kg of formulated IgA2_mRNA_, 5 mg/kg of Sal4 IgA2_R_ or 4.5 mg/kg IgA isolated from human serum. Concentrations of antibody were measured in serum over time by isotype-specific ELISA and modelled using a flexible linear mixed effects model. **(A)** Observed animal-level expression by ELISA and model-based estimates of the mean and 95% credible interval and **(B)** model-based half-life estimates with 95% credible intervals are shown.

### IgA2_mRNA_ has increased sialylation of all asparagine sites as compared to IgA2_R_

IgA2_R_ produced in HEK293 cells has been reported to be under-sialylated compared to IgA isolated from human serum (*23*), resulting in rapid clearance from circulation by the asialoglycoprotein receptor (ASGPR) and other mechanisms (*9, 24*). To determine if glycosylation differences partially explain the more favorable pharmacodynamic properties of IgA_mRNA_, we quantified the levels of sialylation and overall glycosylation patterns of Sal4 IgA2_mRNA_ purified from mouse serum (**Fig. 3**). As serum IgA is mostly monomeric, glycosylation patterns of IgA2_mRNA_ were compared to isolated _m_IgA2_R_ or IgA2_R_ containing monomers and dimers. IgA2_mRNA_ exhibited greater N-glycosylation of the complex type (90%) compared with IgA2_R_ (30%) and _m_IgA2_R_ (19%) (**Fig. S3 and Fig. S4**). In contrast, high-mannose type branched structures were predominant in IgA2_R_ (66%) and _m_IgA2_R_ (76%), as compared to the low levels (5%) on IgA2_mRNA_. The observation of a high level of high-mannose glycosylation of _m_IgA2_R_ and IgA2_R_ correlated with a low level of sialylation (13% - 14%) in these proteins, as compared to the high levels of sialylation (91% of glycan structures in all asparagine sites) of IgA2_mRNA_ (**Fig. 3** and **Fig. S4**). In addition, IgA2_mRNA_ and _m_IgA2_R_ have minimal fucosylation (13-14% of glycan structures at all asparagine sites; **Fig. S4**) compared to IgA2_R_ (24%). High-mannose type branched structures of JC were found to predominate in IgA2_R_ while IgA2_mRNA_ and _m_IgA2_R_ were undetectable (**Fig S5**), further indicating the mostly monomeric nature of IgA2_mRNA_ in circulation.

**Fig. 3.**
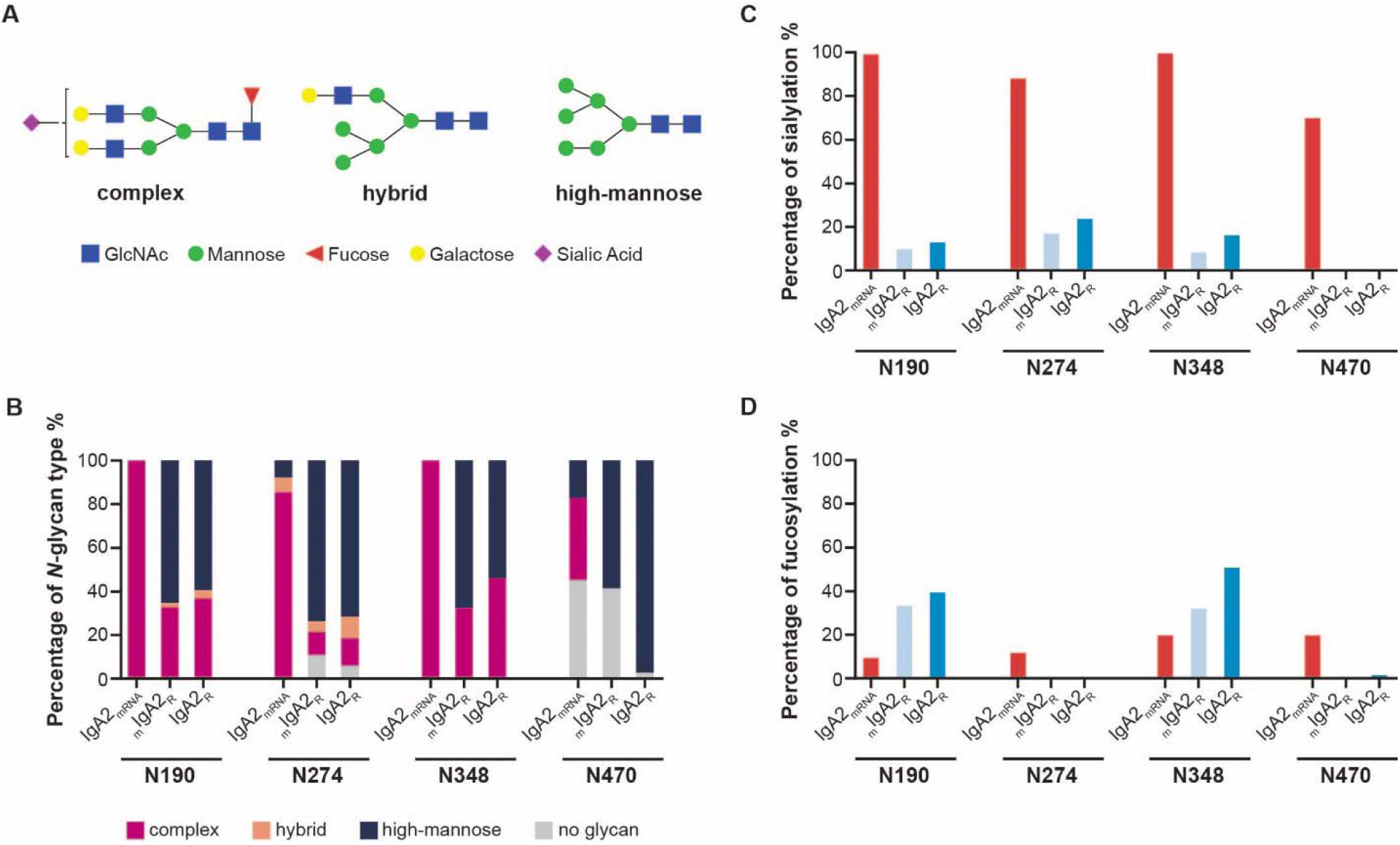
Site-specific N-glycosylation analysis of Sal4 IgA2 heavy chain expressed in mice via mRNA administration or in Expi293 cells via cDNA transfection. **(A)** Examples of three main types of *N*-glycan structure: complex, hybrid, and high-mannose; **(B)** IgA2 *N*-glycan compositions observed at 4 asparagine residues (N190, N274, N348, and N470); **(C)** Sialylation of IgA2 asparagine residues, expressed as a percentage of complex and hybrid glycans that are sialylated; **(D)** Fucosylation of IgA2 asparagine residues, expressed as a percentage of complex, hybrid, and high-mannose glycans that are fucosylated. Consistent detection of glycopeptides was observed across three technical executions in a serum pool from 50 animals.

When *N*-glycosylation was examined at the individual residue level, similar trends in glycosylation patterns were observed (**Fig. 3**). Residues N190 and N348 of IgA2_mRNA_ were found to have nearly 100% complex type glycans. In contrast, N190 and N348 of IgA2_R_ had 31% to 45% complex-type glycans. However, compared to N190 and N348, lower levels of complex type glycans were observed on N274 and N470 in all the IgA2 preparations tested. In addition, N470 was glycosylated at low occupancy (55% for IgA2_mRNA_ and 58% for _m_IgA2_R_) relative to the nearly complete occupancy observed at most other sites.

### IgA2_mRNA_ is detectable in serum and feces, whereas IgG1 is only detectable in serum

Since _d_IgA is naturally transported to mucosal secretions, we investigated whether administration of mRNA/LNP formulated Sal4 IgA2_mRNA_ results in accumulation of _d_IgA_mRNA_ in mucosal tissues. We compared Sal4 _d_IgA2_R_, IgA2_mRNA_ and IgG serum kinetics and mucosal localization over time. Intravenous administration of IgG1_R_ (2.5 mg/kg) to BALB/c mice resulted in immediate systemic circulation of antibodies; however, administration of 2.5mg/kg of _d_IgA2_R_ was undetectable in the serum at any time point examined, potentially due to rapid transport to mucosal tissues (**Fig. 4A**). In contrast, 1 mg/kg of mRNA/LNP encoding for IgG1_mRNA_ and IgA2_mRNA_ reached peak systemic titers of 69 µg/mL and 27 µg/mL, respectively, 24 to 48 hours following injection (**Fig. 4A**).

**Fig. 4.**
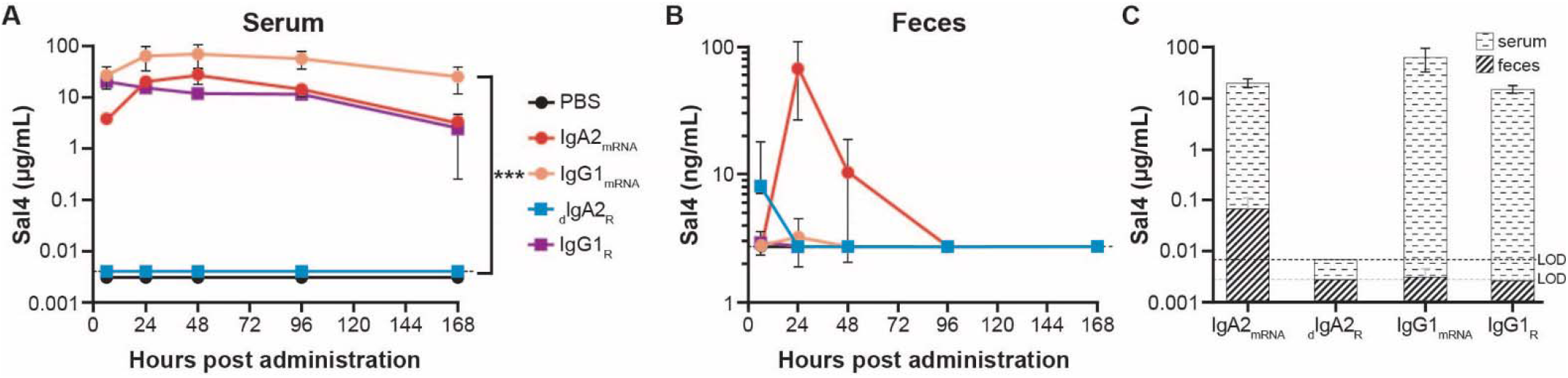
IgA2_mRNA_ expresses in serum and traffics to mucosa. Balb/c mice were injected intravenously with 1 mg/kg of formulated antibody encoded by mRNA modified with 3’idT, 2.5 mg/kg of IgG1_R_ or 2.5 mg/kg _d_IgA2_R_. Concentrations of antibody were measured in **(A)** serum and **(B)** feces over time by isotype-specific ELISA. **(C)** Comparison of serum and feces levels at 24 hours following administration in mice receiving different test articles. Dashed line represents the limit of detection of the assay. Each point represents the mean ± SD of 4-8 mice per group. *** P<0.001 using a One-way ANOVA Kruskal-Wallis test.

Neither IgG1_R_ nor IgG1_mRNA_ were detectable in fecal pellets collected at 24 hours, even though mice had high corresponding concentrations of IgG in sera (**Fig. 4B**). Intravenous injection of Sal4 _d_IgA2_R_ resulted in detectable Sal4 IgA in fecal pellets at 6 hours but it was below the limit of detection for all subsequent time points (**Fig. 4B**). In contrast, *in vivo* expression of IgA2_mRNA_ resulted in mucosal antibody enrichment with an average of 67 ng/mL at 24 hours in fecal pellets (**Fig. 4B, C**, and **Fig. S6**), which remained detectable for 96 hours.

### IgA2_mRNA_ led to a more significant decrease in STm invasion in vivo as compared to IgA2_R_ and IgG

We next used a mouse model of STm intestinal invasion to determine if expression of Sal4 IgA2_mRNA_ was protective in the GI tract (*19*). Mice were administered 1mg/kg of mRNA/LNP encoding for Sal4 IgA2_mRNA_ or IgG_mRNA_ or 5mg/kg analogous recombinant antibody protein. Prior to challenge, serum IgA2_mRNA_ levels were an average of 11 µg/mL, while IgA2_R_ were undetectable (**Fig. 5A, Fig. S7A**). In contrast, IgG1_mRNA_ and IgG1_R_ resulted in significantly higher serum IgG concentrations than IgA, with averages of 291 µg/mL and 84 µg/mL, respectively (**Fig. 5A**). However, in the fecal pellets, IgA2_mRNA_ was found at the highest concentrations (125.2 ng/mL), followed by Sal4 IgG1_mRNA_ (51.66 ng/mL), and IgG1_R_ (9.71 ng/mL) (**Fig. 5B, Fig, S7B**). The elevated levels of fecal IgA2_mRNA_ relative to serum antibody levels is consistent with active transport of Sal4 IgA_mRNA_ into intestinal secretions, while the observed fecal IgG levels are consistent with transudation (**Fig. 5C**). Importantly, IgA2_mRNA_ reduced invasion of intestinal tissues by STm significantly more effectively than either IgA2_R_ or IgG1 (**Fig. 5D**). Protection, as defined by a competitive index (CI) of <0.5, was observed in mice with fecal Sal4 IgA2_mRNA_ levels that were >100 ng/mL, suggesting a threshold of protection associated with mRNA-based delivery (**Fig. S8A-F**).

**Fig. 5.**
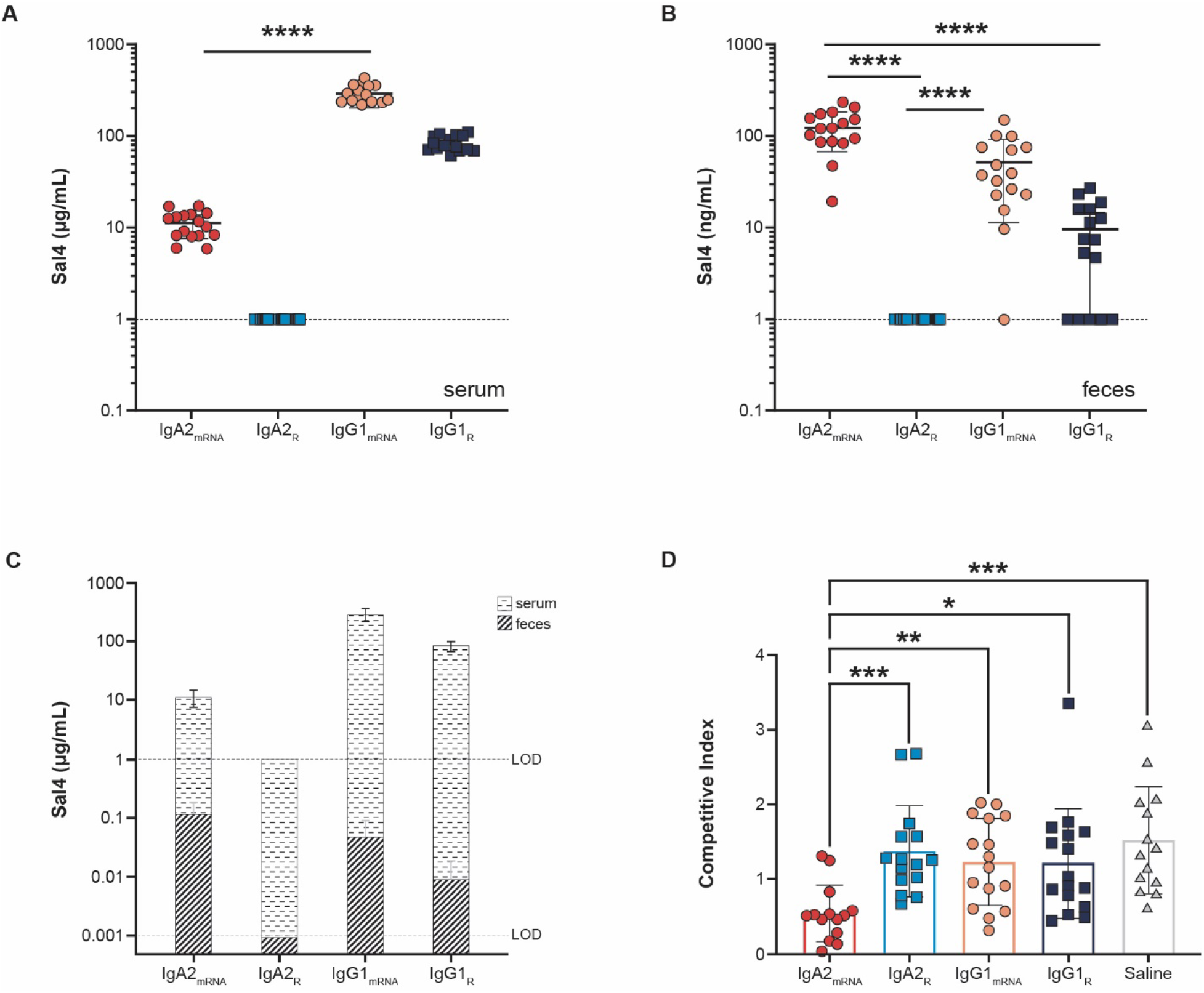
IgA2_mRNA_ blocks STm invasion into mouse Peyer’s patches. Adult BALB/c mice were injected intravenously with 1 mg/kg of formulated mRNA modified with 3’idT and encoding an antibody, 5 mg/kg of recombinant antibody protein or saline. Mice were challenged 24 hours later with an oral gavage of a one-to-one mixture of O5+ and O5-strains (4×10^7^ CFU total). Concentrations of antibody were measured in **(A)** serum and **(B)** feces prior to challenge by isotype-specific ELISA. **(C)** Comparison of serum and fecal antibody levels immediately prior to challenge. **(D)** Competitive indices of O5+ and O5-STm. Shown are the combined results of four independent experiments with four mice per group for a total of 16 mice. Dashed line represents the limit of detection of the assay. Each symbol represents an individual mouse. Statistical significance evaluated for each treatment as compared to IgA2_mRNA_ group, as determined by Kruskal-Wallis test and Dunn’s post-hoc test.

### IgA1_mRNA_ and IgG1_mRNA_ protected mice against lethal challenge in a P. aeruginosa acute pneumonia model

We next investigated the capacity of IgA1_mRNA_ to protect from PA in an acute lethal pneumonia model. CAM003, a well-characterized IgG antibody that binds Psl of PA, was class-switched to IgA1, the most common IgA isotype in the respiratory tract (*25*). mRNA encoded, class-switched CAM003 retained its affinity for PA (**Fig. S9)**, bound to immobilized pIgR and expressed at high levels in mice (**Fig. S9**). Prior to intranasal challenge, adult BALB/c mice administered 1mg/kg mRNA/LNP expressed an average of 68.73 µg/mL IgG1_mRNA_ compared to an average of 7.95 µg/mL of IgA1_mRNA_ in circulation **(Fig. 6A**). Despite significant differences in circulating antibody titers, both IgA1_mRNA_ and IgG1_mRNA_ protected mice from lethal challenge, consistent with the ability of IgA1_mRNA_ to migrate to another mucosal site and demonstrates the ability of mRNA-encoded IgA to protect in this pneumonia challenge model (**Fig. 6B**).

**Figure 6.**
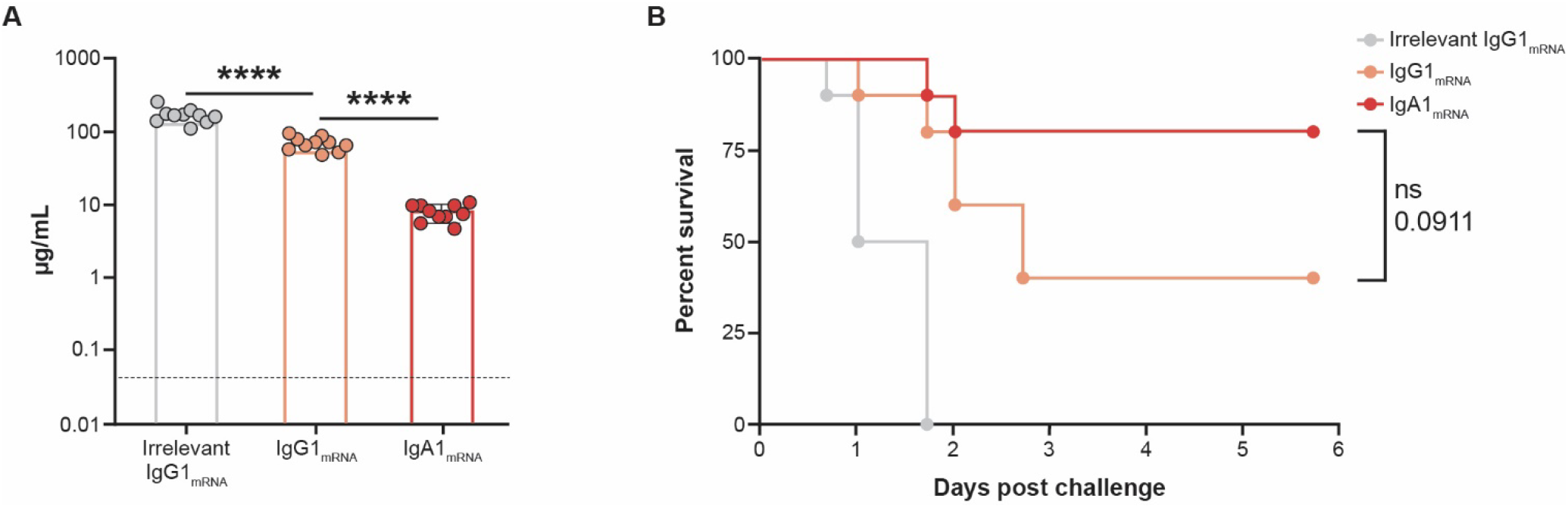
CAM003 IgA1_mRNA_ and IgG1_mRNA_ protects mice against lethal challenge in a *P. aeruginosa* acute pneumonia model. Adult BALB/c mice were injected intravenously with 1 mg/kg of formulated mRNA encoding CAM003 as an IgG1 or an IgA1 and were intranasally challenged 24 hours later with 6.75 log_10_ CFU of *P. aeruginosa* strain PA01. Mice were monitored for survival for 6 days following challenge **(A)** Circulating antibody concentration in serum prior to challenge by isotype-specific ELISA (n=10/group). **(B)** Kaplan-Meier survival curves; differences in survival were calculated by the log-rank test for IgA1 versus IgG1.

## Discussion

As the predominant antibody isoform in mucosal secretions, IgA serves as a first line of defense in immunity to a range of respiratory and enteric pathogens. The compartmentalization of IgA is not by chance; rather, mucosa-derived _d_IgA is actively transported by the pIgR across intact epithelial barriers into external secretions under homeostatic conditions at a rate of 3 to 5 g per day in humans (*26*). This contrasts with IgG, which collects in secretions primarily by transudation, often following inflammation and/or physical breaks in barrier integrity (*27*). The half-life of IgA, once in external fluids, also exceeds that of IgG, due to the presence of the secretory component and glycans that shield IgA from resident proteases (*28*). These unique attributes, along with the demonstrated ability of certain IgA to intercept and incapacitate pathogens prior to accessing mucosal tissues have raised the prospect of deploying IgA mAbs as novel interventions to combat infectious diseases of the gut and upper airways. However, advances on this front have been stymied by the cost of generating uniform human _d_IgA_R_ mAbs at scale.

In this study, we are the first to report on the use of mRNA to encode human IgA mAbs that resemble native IgA in their biochemical, biophysical, and functional properties. In a mouse model, intravenous LNP-based delivery of a mRNA encoding Sal4 IgA2 and J chain resulted in the accumulation of antigen-specific IgA to levels sufficient to impair STm invasion into gut-associated lymphoid tissues. The degree of protection obtained from Sal4 IgA2_mRNA_ was similar to that achieved with purified, recombinant Sal4 SIgA2 administered orally (*29*). Neither Sal4 IgG_mRNA_ or IgG_R_ were protective despite high levels in serum. Thus, our results are consistent with Sal4 IgA2_mRNA_ intercepting STm in the intestinal lumen and blocking uptake into Peyer’s patch tissues. In the case of PA, both IgA1_mRNA_ and IgG1_mRNA_ derivatives of CAM003 were sufficient to limit bacterial colonization of the lung where both isotypes are known to play a role in mucosal immunity.

Another benefit of IgA_mRNA_ is that it displayed pharmacodynamics *in vivo* more like endogenous human IgA than IgA_R_. This observed dichotomy in serum half-life between IgA2_mRNA_ and IgA2_R_ could be due to differences in glycosylation patterns as a result of different cell type expression (*9, 30*). The higher level of sialylation observed for IgA2_mRNA_ is reminiscent of IgA2 isolated from human serum and this could contribute to increased serum half-life via protection from receptor-mediated clearance. Alternatively, the high mannose glycosylation of IgA2_R_ may contribute to faster clearance, which has been observed for IgG (*31*). Lastly, observed differences in fucosylation levels between IgA2_mRNA_ and IgA2_R_ could contribute to differences in half-life (*32*); however, the roles of the IgA2 core fucose in transcytosis and receptor engagement are currently unknown (*33*).

Better pharmacodynamic properties *in vivo* translated to enhanced protection from mucosal bacterial challenge. mRNA/LNPs delivered intravenously resulted in accumulation of levels of antigen-specific IgA in the gut and respiratory tract to attenuate *Salmonella* and *P. aeruginosa* infections. In a stringent model of STm invasion, Sal4 IgA2_mRNA_ resulted in significantly decreased bacterial burden as compared to IgA_R_ and IgG due to its accumulation at the site of infection. Despite significant differences in circulating antibody titers, CAM003 IgA1_mRNA_ protected mice more efficiently from a lethal PA challenge than the analogous IgG_mRNA_, again demonstrating a fundamental advantage of the IgA isotype to accumulate at mucosal sites of infection. While the mucosal localization of IgA may differ significantly between humans and mice due to pIgR localization (*34, 35*), monoclonal IgA produced from mRNA may serve as an important modality in the battle against infectious diseases. The data presented here and in previous studies (*16, 17*) establish mRNA as a central platform for evaluating basic biology and therapeutics in a manner previously unattainable with protein-based approaches. These results illustrate the advantages of mRNA technology for generating protective mucosal mAbs with enhanced activity *in vivo* and provide a path forward for the development of IgA mAbs.

## General

We thank Dr. Gregory Hendricks and Dr. Kyoung Hwan Lee from the Electron Microscopy Facility at UMass Chan Medical School for their support in the use of Talos L120C. At the Wadsworth Center, we thank Dr. Graham Willsey for STm strain construction, Dr. Jenifer Yates for assistance with mouse studies, and the Media and Cell Culture core facility for bacterial growth media. Medical writing and editorial assistance, under the direction of the authors, was provided by Jared Mackenzie, PhD of MEDiSTRAVA in accordance with Good Publication Practice (GPP3) guidelines and funded by Moderna.

## Author Contributions

Conceptualization: CED, AFR, TY, DL, RK, NJM, AC, OJP

Data acquisition: CED, AFR, JED, SKL

Formal analysis: CED, AFR, TY, YL, DL, CJH, ILR, HK, MK, MEP, WJW, JED, SKL

Funding acquisition: CED, AC, OJP

Investigation: CED, AFR, TY, MJM, ZW, YL, DL, MEP,WJW, JED, SKL

Methodology: CED, AFR, MJM, ZW, YL, BRF, SH, EN, DL, RK, SL, HK, MK, MEP, WJW, JED, SKL

Project administration: NJM, AC, OJP

Resources: BRF, SH, EN, SL, MEP, WJW, NJM, AC, OJP

Supervision: YL, RK, SL, NJM, AC, OJP

Validation: AFR, TY, JED, SKL

Visualization: CED, AFR, TY, MJM, ZW, DK, RK, JED, SKL

Writing-original draft: CED, DL

Writing-review & editing: CED, AFR, YL, SH, EN, DL, RK, SL, WJW, NJM, AC, OJP

Approval of final draft: CED, AFR, TY, MJM, ZW, YL, BRF, SH, EN, DL, RK, SL, CJH, ILR, HK, MK, MEP, WJW, JED, SKL, NJM, AC, OJP

## Data and Material Availability

The authors declare that the data supporting the findings of this study are available within this article and its Supplementary Information.

