## Supplementary Materials for "mRNA delivery of dimeric human IgA protects mucosal tissues from bacterial infection"

**This Word file includes:**

**Materials and Methods**

**Table S1.**

**Figs. S1 to S9**

### **Supplementary Material.**

#### **Materials and Methods**

Cell lines

Expi293F cells were obtained from ThermoFisher (A14527) and maintained in Expi293 expression medium (ThermoFisher) in an Infors incubator with 5% CO_2_ shaking at 120 rpm. HeLa cells were obtained from ATCC (Cat. # CCL-2) and maintained in Dulbecco’s Modified Media (DMEM) with 10% fetal bovine serum at 37°C and 5% CO_2_.

Bacterial strains and growth conditions.

*Salmonella enterica* serovar Typhimurium (STm) was purchased from ATCC (ATCC 14028s). *S. Typhimurium* strains GGW445 (*sj8101::kan oafA126::*TN*10d-*Tc *fkpA-lacZ*) and GGW444 (*zj8101::kan*) with the O4 and O5 O-Ag, respectively, are derivatives of ATCC 14028s (*19, 36, 37*). *Pseudomonas aeruginosa* PA01 strain was purchased from ATCC (ATCC 15692).

Recombinant antibody proteins.

Recombinant protein versions of all monoclonal antibodies were purchased from Genscript or GeneArt (ThermoFisher). CAM003 IgG (GeneArt) was purified from transient transfection of EXPI293 with HiTrap Protein A HP and polished using HiLoad Superdex 200 26/600 prep. Sal4 IgG (Genscript) was purified from transient transfection of HD CHO-S cells with MabSelect SuRe^TM^ LX. Sal4 IgA2m1 (GeneArt) was purified from transient transfection of EXPI293 with CaptureSelect^TM^ IgA Affinity matrix. All protein concentrations were determined by absorption at 280nm using NanoDrop.

Generation of modified mRNA and LNPs.

Sequence-optimized mRNA encoding functional IgA monoclonal antibodies were synthesized *in vitro* using an optimized T7 RNA polymerase-mediated transcription reaction with complete replacement of uridine by N1-methyl-pseudouridine (*38*). The reactions included a DNA template containing an open reading frame flanked by 5′ untranslated region (UTR) and 3′ UTR sequences with a terminal encoded polyA tail. Where indicated, inverted deoxythymidine (idT) were appended to the 3´ terminus of fully synthesized mRNA by phosphodiester linkage of a modifying oligo with 3´ idT. Free and end-stabilized mRNA were purified, buffer exchanged and sterile filtered.

Lipid nanoparticle-formulated mRNA was produced through a modified ethanol-drop nanoprecipitation process as described previously (*30*). Briefly, ionizable, structural, helper, and polyethylene glycol lipids were mixed with mRNA in acetate buffe at a pH of 5.0 and at a ratio of 3:1 (lipids:mRNA). The mixture was neutralized with Tris-Cl at a pH 7.5, sucrose was added as a cryoprotectant, and the final solution was sterile filtered. Vials were filled with formulated LNP and stored frozen at –70°C until further use. The drug product underwent analytical characterization, which included the determination of particle size and polydispersity, encapsulation, mRNA purity, osmolality, pH, endotoxin and bioburden, and the material was deemed acceptable for *in vivo* study.

mRNA transfection for mAb expression.

EXPI293 cells were transiently transfected with the corresponding mRNA encoding for Sal4 IgG, Sal4 IgA2m1, CAM003 IgG or CAM003 IgA1 HC, LC and/or JC using Trans IT-mRNA Transfection Kit (Mirus Bio LLC) per the manufacturer’s recommendations. For all IgG isotypes, this required co-transfection of HC and LC whereas all IgA isotypes consisted of HC, LC and JC transfections unless otherwise stated that JC was not included. EXPI293 cells were diluted to 1x10^6^ cells/mL with EXPI293 Expression Medium (ThermoFisher). Using a ratio of 1 μg mRNA per 1 mL of culture, mRNA was first added to Opti-MEM I SFM (Gibco), followed by addition of TransIT-mRNA transfection reagent (Mirus Bio) at a 1:2 ratio of mRNA to TransIT. The complex was allowed to form for 3 minutes before addition to the suspension culture. For DNA transfections, EXPI293 cells were transfected at 3x10^6^ cells/mL using 1 μg plasmid DNA per mL of culture. DNA complexes were formed by separately diluting DNA and Turbo293 Transfection reagent (Speed Biosystems) in Opti-MEM I, then combined and allowed to incubate at RT for 20 minutes before adding to suspension culture. At 48 h following transfection, cell cultures were centrifuged, and supernatant was collected. Unless being purified, majority of supernatant was further concentrated in 50 to 100 kdA Amicon filters for 40 minutes at 3000 x g. Transfection supernatants and concentrated supernatants were further used to quantitate and characterize by enzyme-linked immunosorbent assay (ELISA).

Enzyme-linked immunosorbent assay (ELISA).

To quantitate total human IgG (hIgG) or IgA (hIgA), 96-well NUNC Maxisorp plates (ThermoScientific) or Immulon 4HBX plates (Thermo Fisher Scientific) were coated with 0.1 mL per well of goat anti-human IgG Fc fragment (Bethyl A80-104A) or goat anti-human IgA (Bethyl A80-102A) at 1:100 dilution in 0.05 M carbonate-bicarbonate (Sigma C30411) overnight at 4°C. To determine binding to human pIgR, Nunc maxisorp plates were coated with 2 µg/mL of recombinant human pIgR (R&D Biosystems 2717-PG-05) at 4°C overnight. For both isotype-specific and pIgR ELISA, plates were washed with an automated plate washer (Biotek) or 4x with 0.05% PBS-T and were subsequently blocked with 0.2 mL per well of Superblock PBST (ThermoFisher) or 2% goat serum in PBS-T for 2 hours at room temperature. Using purified antibodies as a standard, mRNA transfection supernatant or mouse serum was serially diluted in PBST in a dilution plate and 0.1mL per well was transferred to the coated plates and incubated for 1 to 2 hours at room temperature. Following incubation, plates were subsequently washed and incubated with 0.1 mL per well of goat anti-human IgG horseradish peroxidase (HRP; Southern Biotech; 1:5000), goat anti-human lambda HRP (Bethyl; 1:10000), goat anti-human kappa HRP (Bethyl; 1:10000) or goat anti-human IgA HRP (Southern Biotech; 1:5000) for 1 hour at room temperature. Plates were subsequently washed and incubated with 0.1 mL per well of Sureblue TMB 1-C substrate (Fisher Scientific) for 5 minutes. The reaction was stopped with 0.1 mL per well of TMB Stop solution (SeraCare) and read at an absorbance of 450nm on a SpectraMax ABS Microplate Reader or a SpectraMax iD3 Microplate Reader (Molecular Devices). Absolute quantities of human antibody in transfection supernatant or mouse serum were extrapolated in GraphPad Prism 9 using a standard curve that was generated with the appropriate purified isotype version of Sal4 or CAM003.

Bacterial whole cell ELISA.

For STm whole cell ELISA (WCE), an overnight culture of STm (ATCC 14028s) was inoculated with one colony from a freshly streaked plate into 5mL of LB broth. The following day, the overnight STm culture was subcultured 1:50 in LB broth and grown to OD_600_ of 0.7 to 0.8, with a final adjusted concentration of OD_600_ 0.7. Cells were washed 2x with sterile PBS and resuspended in the prewash volume. NUNC MaxiSorp or Immulon 4HBX plates (Thermo Fisher Scientific) were coated with 0.1 mL per well of the STm bacterial solution, covered and incubated at 4°C overnight. For *P. aeruginosa* PA01 WCE, 10 to 50 mL of LB broth was inoculated with one colony from a freshly streaked plate. Bacteria were grown to an OD_650_ of ~2.0 and subsequently diluted with LB broth to an OD_650_ of 0.5, followed by an additional 50% dilution with sterile PBS. NUNC MaxiSorp plates were coated with 0.1 mL per well of the PA01 bacterial solution, covered and incubated at 4°C overnight. For STm WCE, wells were washed 4x with 0.5% PBS-T and blocked in either 0.2 mL of Superblock PBST (Thermo Fisher Scientific) or 2% goat serum in PBS-T for 2 hours at room temperature. For PA01 WCE, wells were washed 4x with 0.5% PBST in an automated plate washer (Biotek) and subsequently blocked with 0.2mL Superblock PBST (ThermoFisher) for 2 hours at room temperature. Subsequent ELISA steps are identical to what has been described previously.

HeLa cell invasion assay.

HeLa cells were obtained from the ATCC and maintained in Dulbecco’s Modified Eagle Media (DMEM) with 10% fetal bovine serum at 37°C and 5% CO_2_. The HeLa cell invasion assay was performed as described (*19*). Cells were seeded at 5x10^5^ cells/mL in 96-well plates and grown for 24 hours until 70% to 90% confluent. Prior to invasion assays, cells were washed 3x with serum-free DMEM. Overnight cultures of GGW444 (O5+) and GGW445 (O5-) were diluted 1:50 into LB at 37°C and 220 rpm and adjusted to an OD_600_ of 0.7. Strains were mixed 1:1 and washed 2x by centrifugation (6,000 x *g* for 4 min) and resuspended in PBS with a pH of 7.4. Bacteria were diluted 1:10 in Hanks’ Balanced Salt Solution (HBSS, Sigma-Aldrich); an aliquot was plated on LB agar supplemented with kanamycin (100 µg/mL) and X-gal (40 µg/mL) for subsequent determination of CFUs associated with bacterial input.

For the invasion assay, bacterial mixtures were incubated with 15 µg/mL of Sal4 IgA for 15 minutes at 37°C. Treated bacteria were applied to HeLa cell monolayers and the culture plate was centrifuged at 1,000 x g for 10 minutes. The microtiter plates were then incubated for 60 minutes at 37°C and the cells washed 3x with HBSS and treated with gentamicin (100µg/mL) to eliminate extracellular bacteria. Cells were then washed 3 times with HBSS and lysed with 1% Triton X-100 (in Ca^2+^ and Mg^2+^-free PBS)., The cells were then serially diluted and plated on LB agar containing kanamycin (100µg/mL) and X-gal (40µg/mL), and then incubated overnight at 37°C. The competitive index (CI) [(%strain A recovered/% B strain recovered)/(%strain A inoculated/% strain B inoculated)] was calculated for each treatment group.

Purification of Sal4 IgA.

Purification proceeded identically for both mRNA and DNA transfections. To harvest, suspension cultures were centrifuged at 4,000 rcf for 30 minutes and pellets were discarded. Cell supernatant was filtered through a 0.22 μm vacuum filter followed by gravity chromatography with columns packed with CaptureSelect IgA Affinity Matrix (ThermoFisher). Columns were washed with PBS before eluting in 0.1 M Glycine pH 3 and were then immediately neutralized with 1 M Tris with a pH of 8. Affinity purified IgA was then injected onto a Superdex 200 Increase 10/300 GL gel filtration column (Cytiva) using an AKTA Pure FPLC.

Transmission electron microscopy (TEM) imaging.

For each sample, a ~3.5 μL aliquot was applied to a freshly glow-discharged, carbon-coated copper grid (Electron Microscopy Sciences, CF300-Cu) and allowed to absorb for 30 seconds. After blotting the excess solution, grids were stained with 1% uranyl acetate solution (Electron Microscopy Sciences) for 1 minute and then air-dried. A Thermo Fisher Scientific Talos L120C electron microscope equipped with a 4k x 4k Ceta CMOS Camera was used for data collection. Images were captured at a nominal magnification of 57,000x and 73,000x (pixel size 2.60 Å/pixel and 2.04 Å/pixel, respectively) with a defocus range of ~ -1.5 to -2 μm. Reference-free 2D classification was performed using Relion 4.0.

Expression of mAbs in mice.

All mouse studies were approved by the Animal Care and Use Committee at Moderna. Six- to 8-week old female BALB/c mice (Charles River Laboratories), in groups of 5 to 28 mice each, were injected intravenously with the indicated recombinant protein, mRNA/LNP, or PBS, at the indicated dose in 100 µL. All *in vivo* studies with mRNA/LNP were coformulations of HC and LC for IgG and HC, LC, and JC for IgA. For IgA, this results in a mixture of monomeric, dimeric, and polymeric IgA. Unless otherwise stated, all IgA_R_ injected into mice is a mixture of monomeric, dimeric, and polymeric. Mice were bled via submandibular vein at the indicated time points and serum was isolated for antibody quantification by ELISA. At the final indicated time point, mice were euthanized via CO_2_ asphyxiation and a terminal bleed was collected via cardiac puncture.

Mouse tissue processing for mAb quantification.

For fecal samples, three to four fresh fecal pellets were collected at each of the indicated time points following intravenous administration of test articles. Total weight of the fecal pellets per mouse were recorded and fecal extract buffer (PBS containing 10% normal goat serum [Gibco] and 1 protease inhibitor per 50mL solution [Pierce A32953]) was added to achieve a final concentration of 200 mg/mL. Pellets were vortexed and manually broken until the entire pellet was disrupted. Following centrifugation at 13,000 rpm for 10 minutes at 4°C, supernatants were collected and mAb concentrations were quantified via ELISA.

Where indicated, intestines were harvested at 168 hours after injection following CO_2_ asphyxiation and washed in PBS to remove any mucous or fecal debris. Each mouse intestine (comprising both the small and large intestine) was weighed and homogenized (2x at 5 m/s for 45 s, with a 10 s break between sets) at 2g/mL in homogenization buffer (PBS + 10% normal goat serum) in 2.0 mL homogenizing tubes containing 2.8 mm ceramic beads (Fisherbrand) using the BeadMill homogenizer (Fisherbrand). Eppendorf tubes containing homogenized intestinal tissue were spun at 13,000 rpm for 10 hours at 4°C to obtain debris-free supernatant which was stored for further analysis by ELISA.

Statistical model for IgA half-life modeling.

Since each analyte was measured at different timepoints, we fit the following model to each analyte separately. We estimated the population-level half-life of each analyte using a flexible linear mixed effects model that incorporates a first-order kinetic component for terminal measurements. To do so, we modeled the terminal decay phase of the data using a first-order kinetic model but employed a flexible spline basis to model the earlier phases. Let $y_{ij}$ be the measured analyte concentration for animal $i$ at time $t_{ij}$, where $i=1,\ldots,n$ and $t_{ij}=t_{i1},\ldots,t_{in_{i}}$. To treat the early and terminal phase of the time course differently, we introduce variables $x_{ij}^{0}$ and $x_{ij}^{1}$, defined as

$$x_{ij}^{0}=\left\{ \begin{matrix} t_{ij} & t_{ij}\leq t \\ 0 & t_{ij}>t \end{matrix} \right. x_{ij}^{1}=\left\{ \begin{matrix} 0 & t_{ij}\leq t \\ t_{ij} & t_{ij}>t \end{matrix} \right. (1)$$

where $t$ is some fixed threshold after which the terminal decay phase has begun. Then,

$\text{log}\left( y_{ij} \right)=\left\{ \begin{matrix} \beta_{0}+\beta_{1}x_{ij}^{1}+f\left( x_{ij}^{0} \right)+\gamma_{0i}+\gamma_{1i}x_{ij}^{1}+\epsilon\left( t_{ij} \right) & y_{ij}^{\star}\geq L \\ \text{log}\left( L \right) & y_{ij}^{\star}<L \end{matrix} \right. (2)$

where $y_{ij}^{\star}$ is the concentration of the analyte for animal $i$ at time $t_{ij}$ prior to censoring by the lower limit of quantification (LLOQ), $L$ is the LLOQ of the assay, $t$ is a threshold after which the decay process has entered the terminal phase, and $f$ is a $k$-dimensional thin plate regression spline basis (*39*).

We allow for animal-specific random intercepts and slopes, where the random slopes imply animal-specific deviations about the population-level terminal IgA half-life. Therefore, we have

$$\boldsymbol{\gamma}_{i}=\left( \begin{matrix} \gamma_{0i} \\ \gamma_{1i} \end{matrix} \right)\overset{\mathrm{iid}}{\sim}N\left( \begin{matrix} \left( \begin{matrix} 0 \\ 0 \end{matrix} \right),\left( \begin{matrix} \tau_{0}^{2} & 0 \\ 0 & \tau_{1}^{2} \end{matrix} \right) \end{matrix} \right) (3)$$

We allow residual variance to change with time following a 7-dimensional thin plate spline regression basis, giving $\epsilon(t_{ij})\sim N(0,\sigma_{t_{ij}}^{2})$.

In order to choose the threshold *t*, for each analyte, we selected the latest *t* to allow for ≥3 timepoints where the majority of measurements were above the LLOQ. Since all IgA2_R_ measurements were above the LLOQ, we selected the *t* such that the last 3 timepoints were used to estimate the terminal half-life. For IgA2_mRNA_ and human serum IgA, the last 4 time points were used to estimate terminal half-life. The dimension of the thin plate regression spline basis *f* was chosen to be as large as possible, given the threshold *t*.

All modeling was implemented using the R package brms (version 2.17.3) (*40*) using default, non- or weakly-informative prior specifications.

Purification of IgA2 from mouse serum.

50 Balb/c mice per group (8-10 weeks old) were intravenously administered either1 mg/kg of the indicated mRNA/LNP formulations or PBS and terminal bleed serum was collected 24 hours following administration (**Fig. S3**). Sal4 IgA2 was purified from pooled mouse sera using peptide M agarose (Invivogen) following the manufacturer’s instructions. Briefly, the sera were concentrated by 100kDa Pierce Protein Concentrators PES (Thermo Fisher), then buffer exchanged with PBS buffer. The sera was then incubated with peptide M agarose resin overnight at 4°C, and the resin was washed using PBS buffer. After another wash step with 0.1 M glycine buffer (pH 5.0), IgA2 was eluted with 0.2 M glycine buffer (pH 2.5) and incubated for 5 minutes at room temperature. The elution step was repeated 3x, and the eluted IgA2 concentration of the administered group and control group were measured by ELISA.

Isolation of recombinant IgA2 monomer.

Sal4 IgA2 expressed by EXPI293 cell line as described above was further using the AKTA pure protein purification system. The monomeric IgA2 (_m_IgA2_R_) was isolated for glycosylation comparison with mRNA expressed IgA2 *in vivo* (**Fig. S3**). Briefly, 1mg of the total IgA2 was injected to Superdex 200 Increase 10/300 GL column (Cytiva) and then IgA2 was eluted by PBS buffer at a constant flow rate of 0.25 mL/min. The eluent was collected by AKTA fraction collector F9-C (0.5 ml per fraction). The fractions of monomer were determined using western blot and combined for the next step of the experiment.

Protein digestion.

Sal4 IgA2 proteins from different sources were concentrated to 20 µL by 100 kDa Pierce Protein Concentrators PES, then buffer exchanged with UA buffer (8 M urea in 50 mM Tris-HCl buffer pH 7.8). Dithiothreitol (Sigma-Aldrich) was added to the solution to a concentration of 10 mM; the solution was then heated at 95°C for 10 minutes. Once cooled to room temperature, iodoacetamide (Sigma-Aldrich) was added to the solution to a concentration of 20 mM. The solution was then incubated in darkness at room temperature for 40 minutes and another 10 mM dithiothreitol was added to quench the reaction. The solution was then diluted with 200 μL of 0.5 mM Glu-Glu (Sigma-Aldrich) in 50 mM Tris-HCl buffer at a pH of 7.8. Endoproteinase GluC (New England Biolabs) was added to make GluC and IgA2 at a 1:10 ratio (w/w), and the solution was incubated at 37°C overnight. The sample was further digested with Trypsin to a final protease:protein ratio of 1:50 (w/w) for 4 hours at 37°C. The reaction was quenched by adding 0.1% Formic Acid (Fisher Scientific) and the digested peptides were desalted using HyperSep C18 Cartridges (Fisher Scientific). The desalted peptides were then dried by Savant SpeedVac Concentrator (Thermo Fisher Scientific) and stored at –80°C.

N-glycosylation site analysis.

*N-*linked glycosylation sites of Sal4 IgA2 were de-glycosylated and labeled with ^18^O using a previously reported method (**Fig. S3**)(*41*). Sal4 IgA2 proteins from different sources were dissolved in 30 µL of 20 mM ammonium bicarbonate (Sigma-Aldrich) in H_2_^18^O. One unit of PNGase F (Sigma-Aldrich) was then added, and the solution was incubated for 1 hour at 37°C to label the *N*-glycosylation sites. The solution was stored at –80°C for mass spectrometry analysis (**Fig. S3**).

Glycopeptide enrichment.

*N-*linked intact glycopeptides of Sal4 IgA2 were enriched by HILIC using a previously reported method (*42*). ZIC-HILIC particles (The Nest Group, Inc.) were packed in C4 HyperSep tips (Thermo Fisher Scientific). The HILIC tip was then conditioned 3x in LC-MS grade water (Fisher Scientific) followed by conditioning (3x) in 1 mL of 80% acetonitrile (ACN), 0.1% trifluoroacetic acid (TFA). Sal4 IgA2 proteins from different sources were dissolved 3x in 80% ACN and 0.1%TFA and loaded onto HILIC tips. The HILIC tips were then washed 3x with 80% ACN and 0.1%TFA, and the glycopeptides were eluted 3x using 100% water, 0.1%TFA. The eluent was dried by lyophilizer (Labconco) and store at –80°C for mass spectrometry analysis (**Fig. S3**).

Mass spectrometry analysis.

Peptides or glycopeptides were dissolved in different volumes of 0.1% formic acid (FA) and separated (in three technical executions) through a Waters ACQUITY UPLC M-Class liquid chromatography system equipped with an Easy-Spray PepMap Neo C18 75 µm X 500 mm column (Thermo Fisher Scientific) coupled to a 20 mm nanoEase C18 Trap Column trap column (Water Corporation). The mobile phase flow rate was 0.3 μL/min with 0.1% FA in water (A) and 0.1% FA 100% acetonitrile (B). The gradient profile was set as the following: 1% to 35% B for 80 minutes, 35% to 60% B for 10 minutes, 60% to 95% B for 10 minutes, 95% B for 5 minutes, and equilibrated in 1% B for 10 minutes. MS analysis was performed using a Thermo Orbitrap Fusion mass spectrometer (Thermo Fisher Scientific). The spray voltage was set at 1.9 kV. Spectra (maximum IT of 100 ms) were collected from 400 to 2000 m/z at a resolution of 120 K followed by data-dependent HCD MS/MS (at a resolution of 6K, stepped NCE 20,30,40, intensity threshold of 1 × 10^5^ and maximum IT of 100 ms) of the 20 most abundant ions using an isolation window of 2 m/z. Charge-state screening was set to only include ions with more than one and less than eight charges.

Data analysis.

*N*-glycosylation sites data from the ^18^O label experiment was processed with Proteome Discoverer 2.5 (Thermo Fisher Scientific). Glycopeptides were identified by Byonic software (Protein Metrics Inc.). The LC-MS/MS spectra of combined GluC/tryptic digests of glycoproteins were searched against the FASTA sequence of Sal4 IgA2 by choosing RKDE as cleavage sites with a maximum of two missed cleavage sites. MS/MS spectra were matched with a tolerance of 10 ppm on precursor mass and 50 ppm on a fragment mass. All protein identification hits had an FDR rate ≤1%. Carbamidomethylation was set as fixed modification, oxidation of methionine, and deamidation of asparagine and glutamine were used as variable modifications. In Proteome Discoverer, ^18^O labeling of Asn (Δm = 2.9848) was set as variable modification to determine *N*-glycosylation sites and *N*-glycan occupancy (**Fig. S3**). Intact-glycopeptide data were searched against the common mammalian *N*-glycans database in Byonic. Glycans were manually categorized according to their composition using the following criteria: HexNAc(2)Hex(9−4) was classified as high-mannose. HexNAc(3)Hex(5−6)X was classified as hybrid (X can be fucose or sialic acid). Other compositions were classified as complex-type glycans. If any of the compositions had a fucose, it was assigned as a fucosylated glycan. Any glycan containing at least one sialic acid was counted as sialylated (*43*). Glycopeptides with more than five MS/MS spectral identified (PSM#) were used for quantifying the percentage of glycans belonging to any of the three categories, which include complex, hybrid, or high-mannose (**Fig. S3**) (*44*). The same method was used to determine the percentage of fucosylation and sialylation among all glycans. IgA2 was undetectable in both HILIC-enriched and ^18^O-labeled aliquots of the serum pool from PBS-administered animals.

STm intragastric challenge.

Female Balb/c mice aged 8 to 12 weeks were obtained from Taconic Biosciences and cared for by the Wadsworth Center Animal Core Facility. Experiments were performed in accordance with protocols approved by the Wadsworth Center’s IACUC. Four mice per group were used and the challenge was repeated four separate times (16 mice in total per group). The challenge experiments were performed as previously described with some modifications (*19*). This model utilizes a competitive infection assay to normalize natural variations in challenge inoculum using two STm strains premixed prior to oral delivery: one expressing the Sal4 epitope, the O5 antigen (O5+), and an O5-null strain (O5-) that expresses ß-galactosidase to easily distinguish the two strains. A reduction in competitive index (CI) only occurs when Sal4 reduces the number of O5+ in mouse Peyer’s patches.

Briefly, mice were injected intravenously with saline, 1mg/kg of formulated mRNA modified with 3’idT, or 5mg/kg of recombinant protein, 24 hours prior to bacterial challenge. Overnight cultures of GGW444 (O5+) and GGW445 (O5-) were subcultured to an OD_600_ of 0.7, combined 1:1 (v/v) and resuspended in sterile PBS pH 7.4. Following this, an aliquot was plated on LB agar containing kanamycin (100 µg/mL) and X-gal (40 µg/mL) at the start of the experiment. Mice were given an STm gavage (~4x10^7^ CFUs in 200 µLs) and sacrificed (CO_2_ asphyxiation followed by cervical dislocation) 24 hours after challenge, following which the small intestine was removed above the cecum for each mouse. The Peyer’s Patches from each mouse were then pooled in 1 mL sterile PBS and placed on ice. Samples were then homogenized 3x for 30 seconds each using a Bead Mill 4 Homogenizer (Fisher Scientific) and the homogenates serially diluted and plated on LB agar (containing 100 µg/mL kanamycin and 40 µg/mL X-gal) and incubated overnight at 37°C. The number of blue and white colonies were determined and the competitive index (CI) was calculated as [(%strain A recovered/% B strain recovered)/(%strain A inoculated/% strain B inoculated)]. Whole plate dilutions at 100 µL per plate were required in order to observe the required number of colonies to calculate CI. Samples containing fewer than 30 CFUs/100 µL were removed and considered “too few to count”.

*P. aeruginosa* lethal acute pneumonia mouse model.

All mouse studies were approved by the Animal Care and Use Committee at the University of North Texas Health Science Center at Fort Worth and conducted according to IACUC protocol 2019-0030. Female Balb/c mice (22 ± 2g) (Envigo Laboratories), 6-8 weeks old, were administered 1 mg/kg (20 µg/mouse in 100 µL) of the indicated mRNA/LNP formulations intravenously (n=8-10/group) and serum was collected 24 hours later to quantitate circulating antibody titers prior to challenge. Mice were subsequently challenged intranasally with 6.75 log_10_ CFU of *P. aeruginosa* PAO1 and monitored for mortality for 6 days. Results were graphed in a Kaplan-Meier survival curve.

#### **Table S1.**

| Abbreviations used: | Description |
| --- | --- |
| _m_IgA | Monomeric IgA – referred to in results when monomeric portion has been isolated and used |
| _d_IgA | Dimeric IgA – referred to in results when dimeric portion has been isolated and used |
| IgA_mRNA_ | Resultant IgA protein made from mRNA that, unless otherwise stated to be without JC, will consist of monomer, dimer and polymers. |
| IgG_mRNA_ | Resultant IgG protein made from mRNA |
| IgA_R_ | Resultant IgA protein made from traditional recombinant production and purification means involving transfection of DNA into EXPI293 cells and purifying protein. Unless otherwise indicated, these protein preparations will include monomer, dimer and polymer |
| _m_IgA_R_ | Isolated monomeric portion of IgA protein made from traditional recombinant production and purification means. |
| _d_IgA_R_ | Isolated dimeric portion of IgA protein made from traditional recombinant production and purification means |
| IgA2 | Referring to IgA of the IgA2 isotype |
| IgA1 | Referring to IgA of the IgA1 isotype |

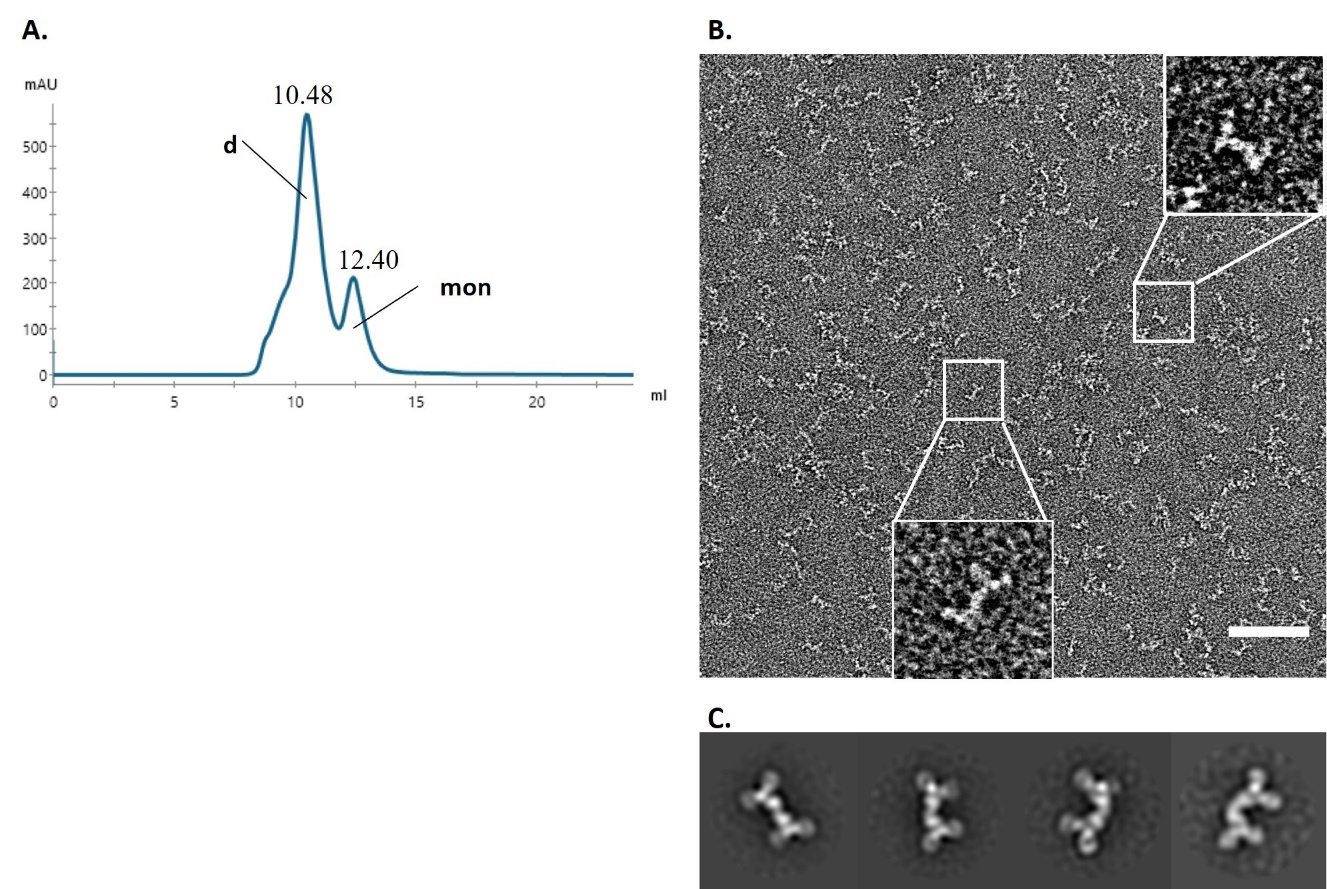

#### **Fig. S1. Purified Sal4 IgA2_R_ made from plasmid transfection of EXPI293. (A)** Analytical size exclusion chromatogram of affinity-purified recombinant IgA2 (IgA2_R_) from transient transfection. Dimer (d) and monomer (mon) peaks are denoted. **(B)** Representative images of negative stain EM of the dimeric peak. Scale bar: 50nm. **(C)** Reference free 2D class averages of dimers.

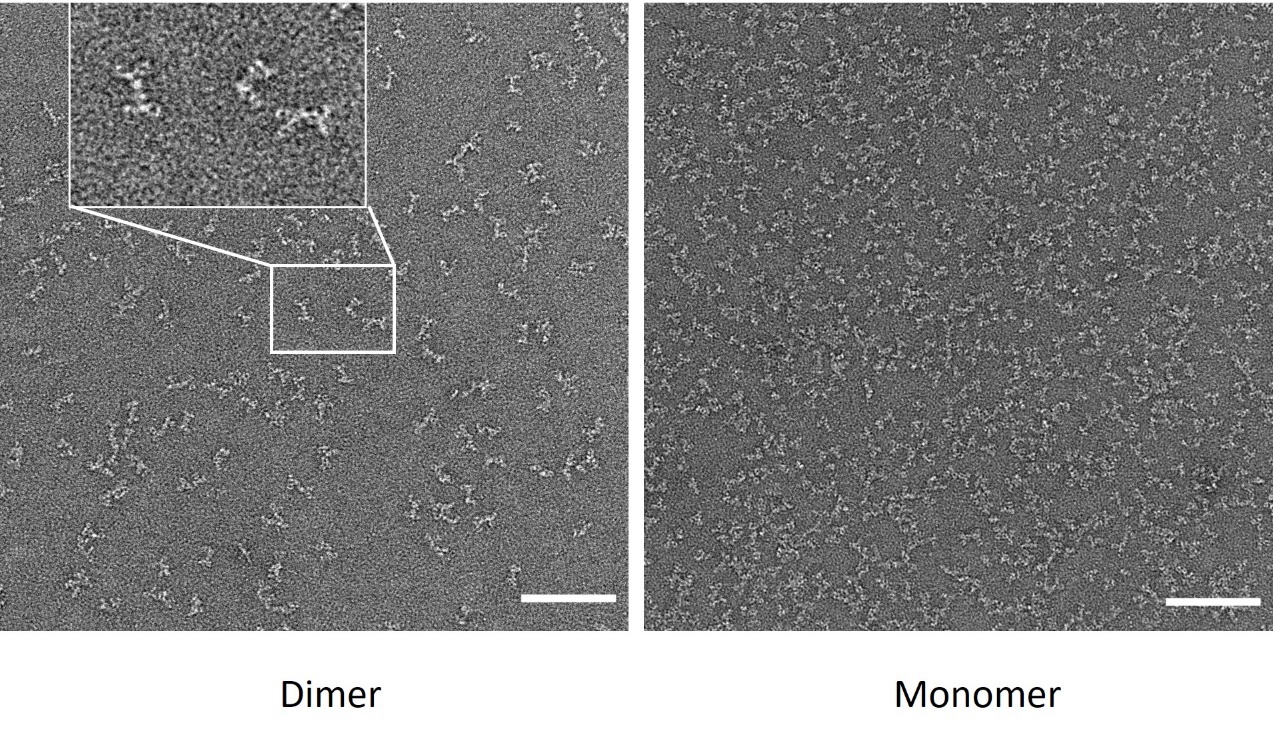

#### **Fig. S2. Raw negative-stain EM (nsEM) images of IgA2mRNA dimer and monomer.** Scale bar: 50nm.

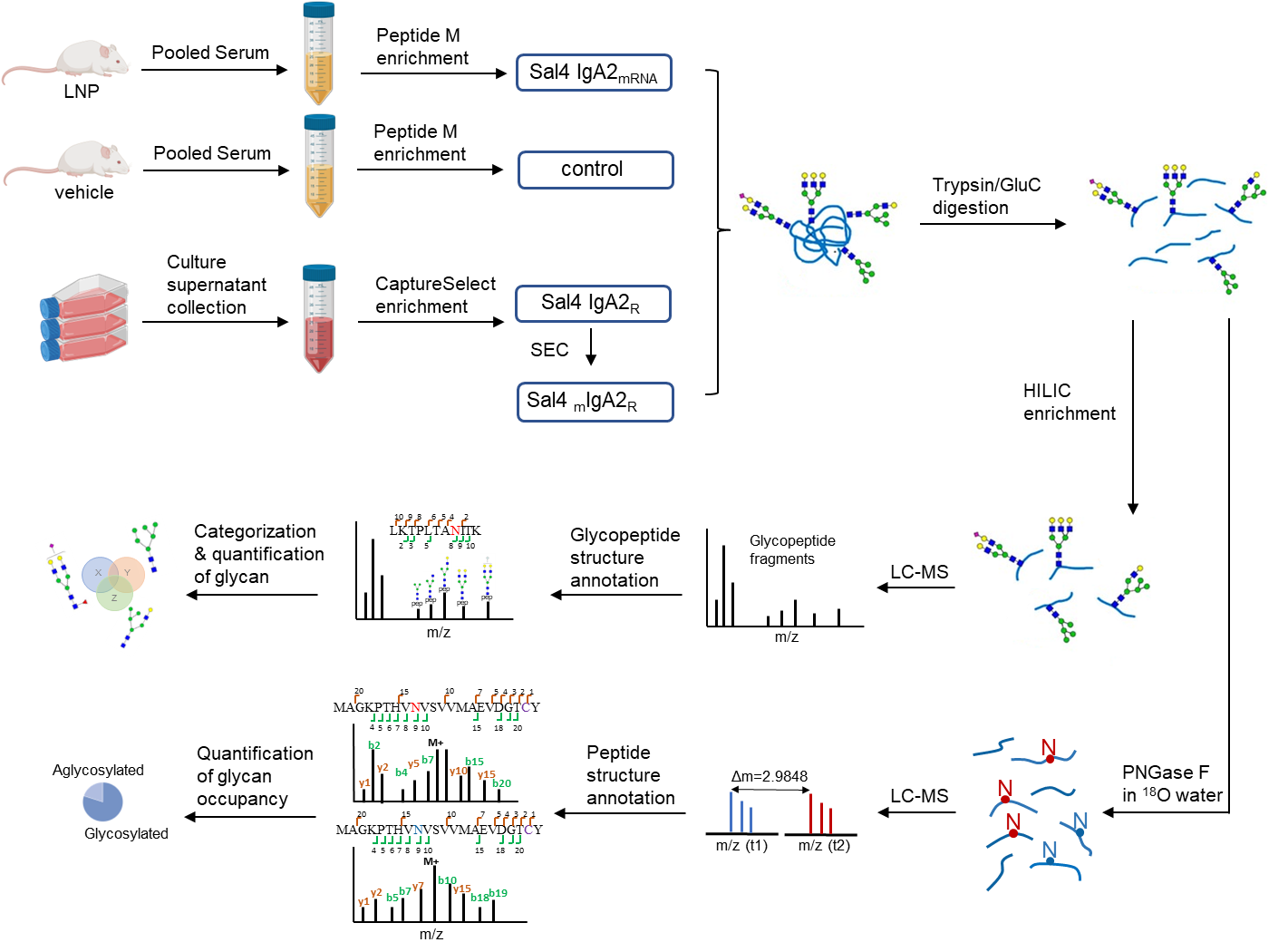

#### **Fig. S3. Workflow of site-specific N-glycosylation analysis of Sal4 IgA2 expressed in mice via mRNA administration (IgA2mRNA) or in EXPI293 cells via cDNA transfection (IgA2R).** 50 Balb/c mice were treated with formulated mRNA encoding Sal4 IgA2, serum was collected after 24 h, and peptide M enrichment was conducted to purify Sal4 IgA2mRNA. The control group was treated under the same conditions but without mRNA formulated in the vehicle. Sal4 IgA2R was synthesized by GeneArt (ThermoFisher) by transfecting EXPI293 cells with cDNA encoding Sal4 IgA2. Cell culture supernatant was collected and both monomeric and dimeric recombinant antibody was purified using CaptureSelect affinity chromatography column (ThermoFisher). To further isolate Sal4 mIgA2R, SEC separation was performed on Sal4 IgA2R. After isolation, both proteins were digested to obtain a peptide and glycopeptide mixture. The 18O label method was used to determine the glycan occupancy of each glycosylation site, and HILIC was applied to enrich glycopeptides. Following data acquisition in LC-MS, data analysis was performed to quantify the glycosylation of IgA2.

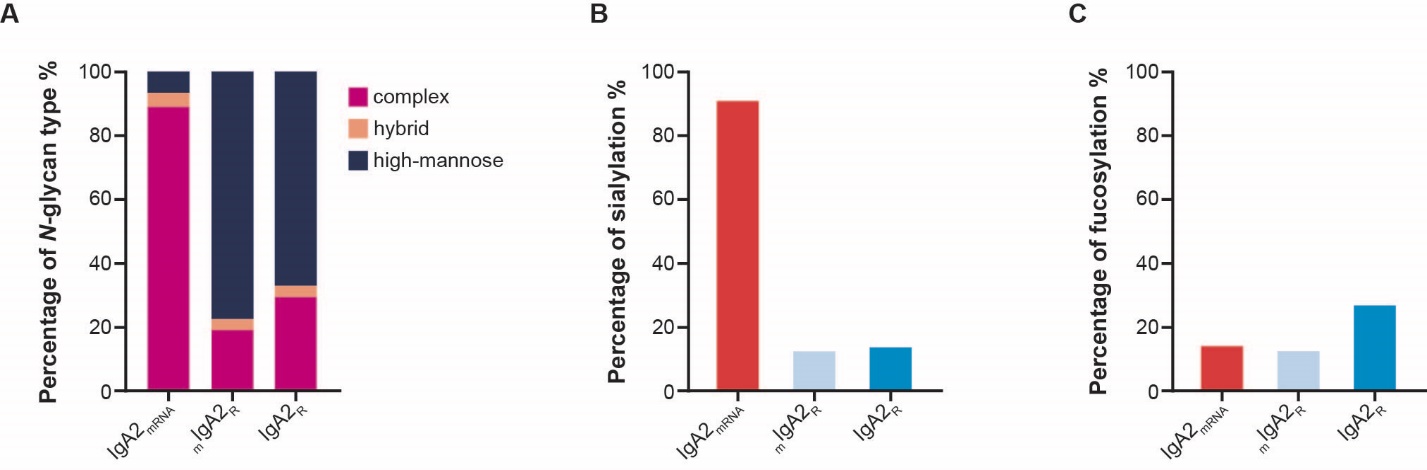

#### **Fig. S4. N-glycosylation analysis of Sal4 IgA2 heavy chain and J chain expressed in mice via mRNA administration or in EXIP293 cells via cDNA transfection.** Glycosylation was measured as the sum of glycosylation levels at four asparagine residues in the heavy chain (N190, N274, N348, and N470) and one in the J chain (N68). (**A**) IgA2 N-glycan compositions; (**B**) Sialylation of IgA2nasparagine residues, expressed as a percentage of complex and hybrid glycans that are sialylated; (**C**) Fucosylation of IgA2 asparagine residues, expressed as a percentage of complex, hybrid, and high-mannose glycans that are fucosylated.

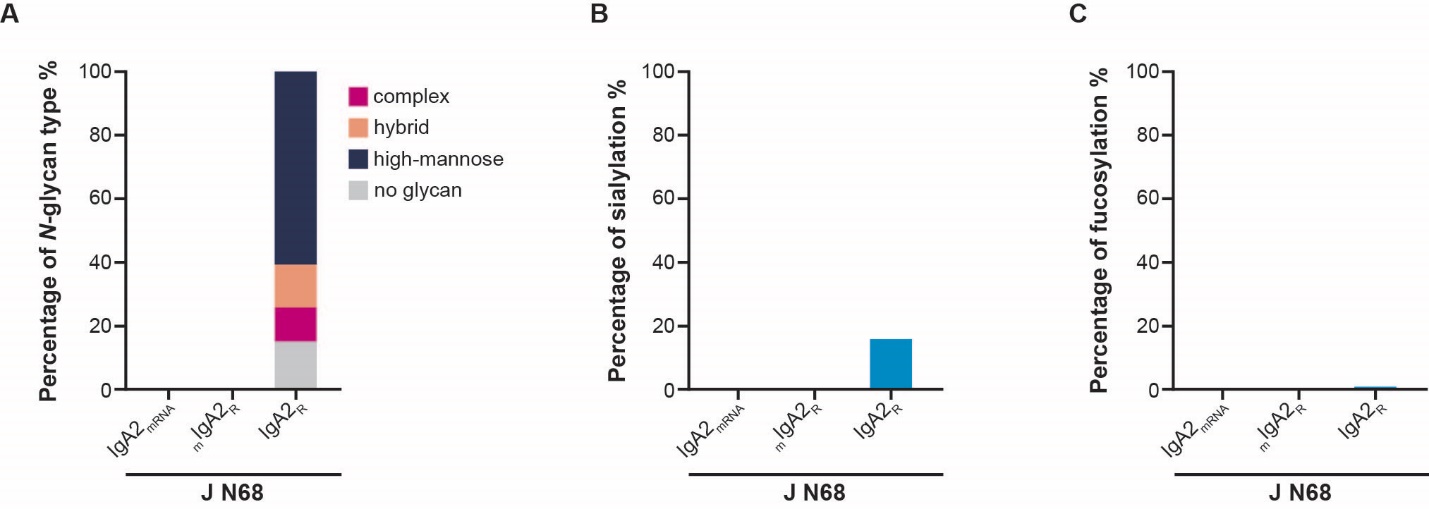

#### **Fig. S5. *N-glycosylation analysis of Sal4 IgA2 J chain (N68) expressed in mice via mRNA administration or in EXPI293 cells via cDNA transfection*. (A)** IgA2 J chain N-glycan compositions at N68; **(B)** Sialylation of IgA2 J chain at N68, expressed as a percentage of complex and hybrid glycans that are sialylated; **(C)** Fucosylation of IgA2 J chain at N68, expressed as a percentage of complex, hybrid, and high-mannose glycans that are fucosylated.

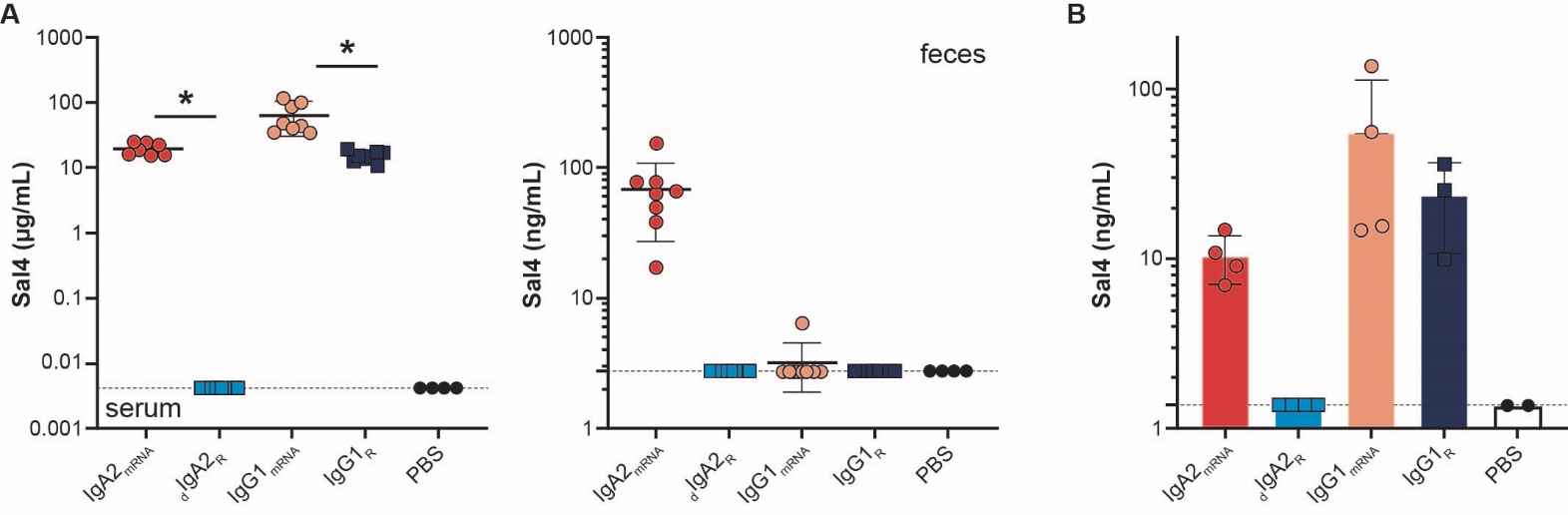

#### **Fig. S6. IgA2_mRNA_ expresses in serum and traffics to mucosa.** Balb/c mice were injected intravenously with 1 mg/kg of formulated mRNA encoded antibody, 2.5 mg/kg of IgG1_R_ or 2.5 mg/kg of _d_IgA2_R_. **(A)** Concentrations of antibody were in serum (left) and feces (right) at 24 hours post injection or **(B)** homogenized intestinal tissue at 168 hours post injection. * P< 0.1 with one-way ANOVA Kruskal-Wallis test

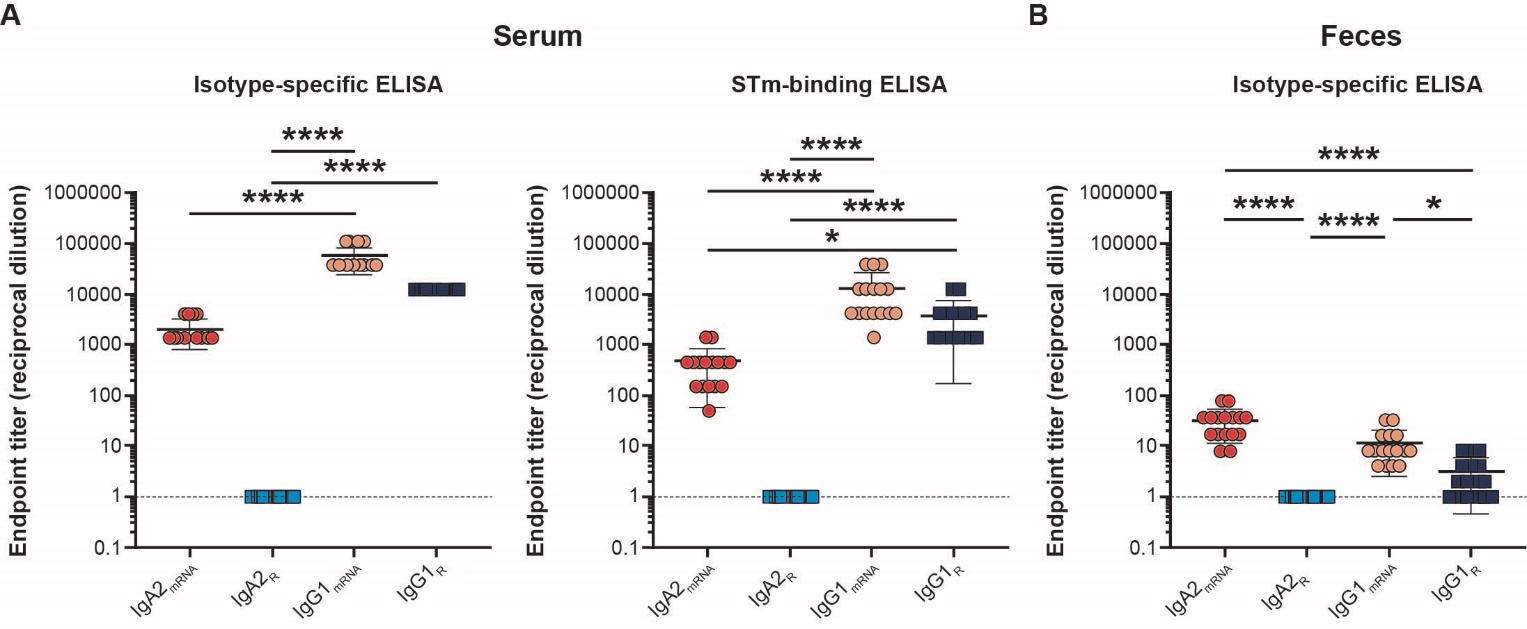

#### **Fig. S7.** Quantitation of antibody in **(A)** serum or **(B)** feces prior to oral challenge with STm reported as endpoint titer (reciprocal dilution). Shown are the combined results of four independent experiments with 4 mice per group for a total of 16 mice. Dashed line represents the limit of detection of the assay. Each symbol represents an individual mouse.

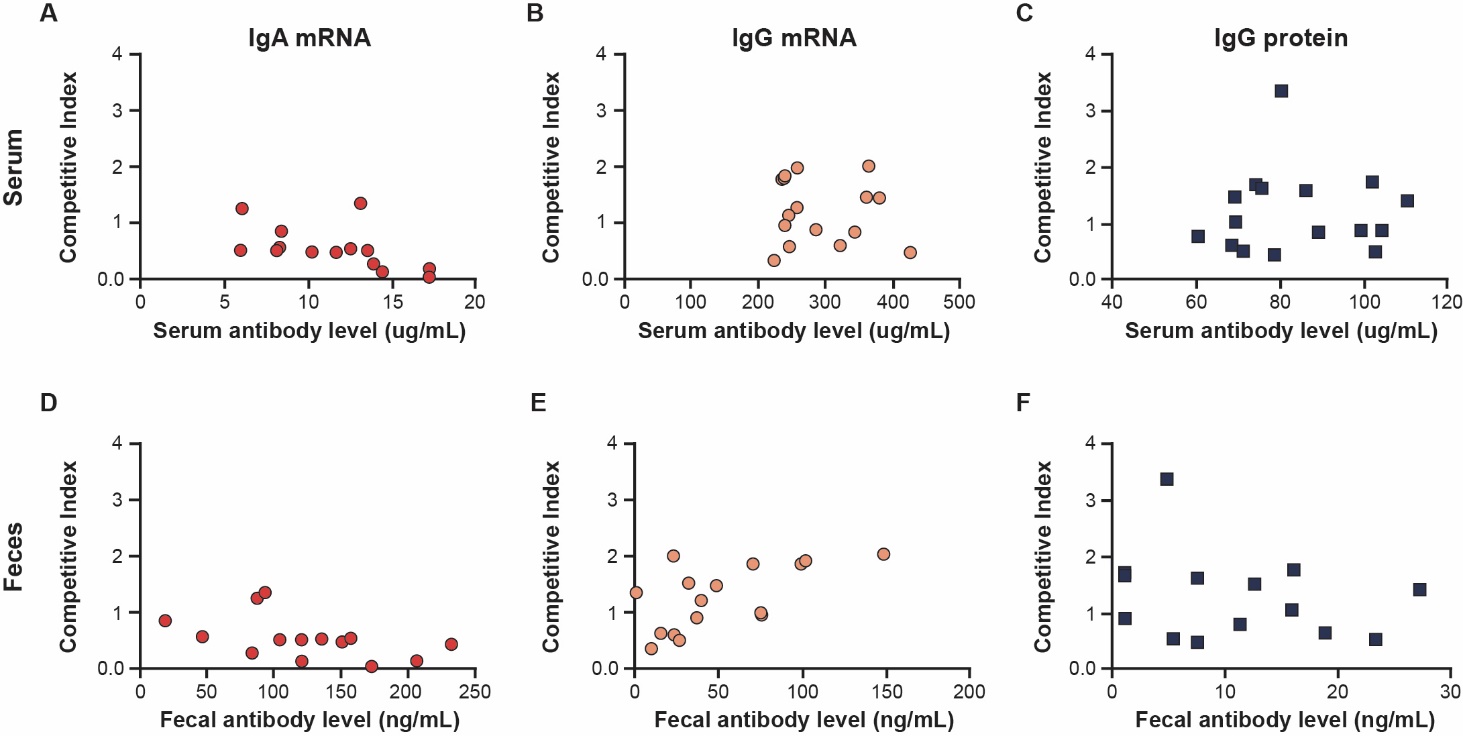

#### **Fig. S8.** Correlations of circulating antibody levels to competitive index in **(A-C)** serum and **(D-F)** feces for **(A, D)** IgA2_mRNA_, **(B, E)** IgG_mRNA_ and **(C, F)** IgG_R_

#### **Fig. S9. IgG1_mRNA_ and IgA1_mRNA_ from in vitro transfection binds to *P. aeruginosa* (PA), pIgR and expresses *in vivo*. (A)** Binding of mRNA transfection supernatant to PA of CAM003 as an IgG or IgA1 isotype. **(B)** Binding of mRNA transfection supernatant of CAM003 IgA1 to human pIgR. **(C)** C57Bl/6 mice were injected intravenously with 0.5 mg/kg of formulated IgG1_mRNA_ or a IgA1_mRNA_. Concentrations of antibody were measured in serum over time by isotype-specific ELISA.

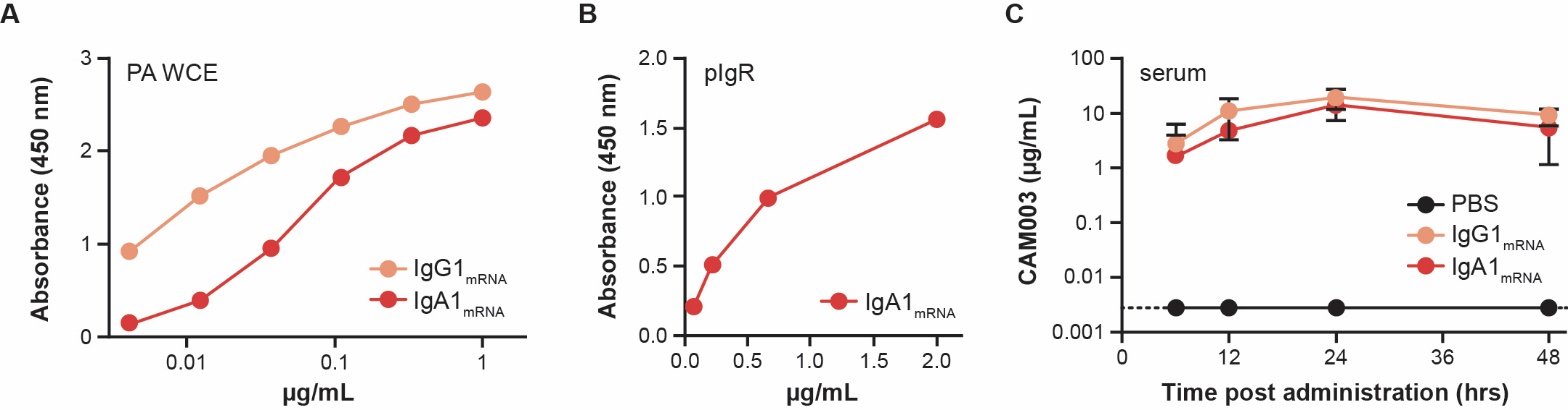
